# Context-dependent ATP7 Interactions with Parkinson’s Disease-associated Genes Modulate Copper Homeostasis Phenotypes

**DOI:** 10.64898/2026.02.02.703386

**Authors:** Brooke M. Allen, Nadia Gonzalez, Erica Werner, Victor Faundez, Alysia D. Vrailas-Mortimer

**Author notes:** Corresponding author: Alysia D. Vrailas-Mortimer, Oregon State University, Department of Biochemistry and Biophysics, Corvallis, OR 97331. Denotes equal contribution (Brooke M. Allen, Nadia Gonzalez).

## Abstract

Copper is an essential micronutrient required by enzymes that catalyze oxygen-dependent reactions, but toxic in excess. Mutations in the ATP7A and ATP7B copper transporters cause neuropsychiatric symptoms and neurodegeneration by mechanisms that remain to be elucidated. We previously reported that the ATP7A biochemical interactome is enriched in Parkinson’s disease (PD) and neurodegeneration associated proteins, yet the functional outcomes of these interactions are unknown. Using *Drosophila*, we tested genetic interactions between ATP7 mutants that alter copper levels and a subset of these PD and neurodegeneration causative genes and found sex differences with some candidate genes enhancing ATP7 deleterious phenotypes in both sexes, while others were sex specific. Most notably, we found that Lrrk2 (Lrrk), the most commonly mutated gene in familial forms of PD, protects against ATP7 dysfunction in epidermal epithelial cells with a stronger effect in males than females. However, in dopaminergic neurons Lrrk plays a role in intracellular copper induced toxicity in females but not males, supporting context dependent interactions between ATP7A and PD-associated genes to protect against disruptions in copper homeostasis.

**Summary Statement:** We performed a genetic interaction screen to explore the relationship between copper homeostasis and Parkinson’s disease and other neurodegeneration associated genes.

## Introduction

Copper is an essential micronutrient that acts as a structural component and catalytic cofactor for key cellular processes, such as antioxidant defense, mitochondrial respiration, energy and iron metabolism, and neurotransmitter production^1–3^. Disruptions in copper homeostasis can cause oxidative damage and impaired function of critical cuproenzymes, resulting in cellular dysfunction and cell death^4–6^. Therefore, copper homeostasis is tightly regulated within the cell by the copper transporters ATP7A and B which pump cytoplasmic copper into the lumen of the Golgi apparatus for metalation of secreted cuproenzymes. Excess levels of cytoplasmic copper, cause ATP7A/B translocation to the plasma membrane, pumping excess copper out of the cell^7– 10^. Thus, deficiencies in ATP7B leads to copper accumulation at the cellular level and at the organism level, resulting in the neurodegenerative disorder Wilson’s disease^5,6^. Whereas ATP7A mutations can cause Menkes disease, a multi-system disorder that includes widespread neurodegeneration caused by decreased copper in the brain due to defective dietary copper absorption^10,11^. Similarly, in the fruit fly, *Drosophila melanogaster*, genetic manipulation of the sole homologue *ATP7* causes a copper depletion phenotype affecting neuronal function^12^ and structure^13^ and induces early mortality^14^. We have found that ATP7A interacts with proteins associated with neurodegeneration and neurodevelopmental disorders^13^, and ATP7A mutant fibroblasts have altered expression of proteins associated with Parkinson’s disease (PD) or neurodegeneration, including elevated expression of the early onset neurodegenerative disease protein, UCHL1^14^. Furthermore, we confirmed this link between ATP7 and UCHL1 through genetic interactions in *Drosophila*^14^.

PD is the second most common neurodegenerative disease and is characterized by loss of dopaminergic neurons in the substantia nigra, impaired mitochondria, and increased oxidative stress^15,16^. Familial forms of PD account for only 5-15% of cases, suggesting a strong environmental component for this disease^15–17^. However, low brain copper levels can lead to dysfunction and neurodegeneration as well, and copper levels are reduced in mice treated with MPTP, a potent inducer of PD-like phenotypes^18,19^. Additionally, key mitochondrial proteins such as Cytochrome c oxidase and superoxide dismutase (SOD1) are cuproenzymes, and ATP7A mutant fibroblasts have impaired mitochondria^14,20^. Furthermore, both elevated and reduced levels of copper can induce oxidative stress through different mechanisms^21,22^, suggesting that both increased and decreased levels of intracellular copper may contribute to PD. This is supported by findings that administering CuSO_4_ or Cu(II)(atsm) improve PD-like phenotypes in mouse models^23–25^, while other studies found improvement with treatment with a copper chelator^26,27^. Furthermore, copper interacts with proteins mutated in familial forms of PD. DJ-1 can act as a copper chaperone to protect against metal toxicity^28,29^, while copper binding to alpha-synuclein promotes its aggregation^30,31^. In addition, loss of ATP7A exacerbated motor phenotypes in a mouse alpha-synuclein (*SCNA*) model^32^, and in *C. elegans*, PARKIN and LRRK2 PD-like phenotypes were enhanced by environmental exposure to copper^33^. These data suggest a strong connection between copper homeostasis and PD, however the mechanism remains poorly understood.

To further explore this link between copper homeostasis and PD, we tested the functional consequences of PD- or neurodegeneration associated genes that we identified as ATP7A/B interactors in our previous proteomics screens^13,14^ genetically interact with ATP7 using the *Drosophila* model system, which we have shown recapitulates many of the phenotypes of ATP7A/B human cell culture models^13,14,20,34^. We used two paradigms previously validated that allow us to differentiate roles of copper homeostasis in non-neuronal cells (the epidermal epithelium) and neuronal cells (dopaminergic neurons) in response to increased cellular copper by ATP7 knockdown (RNAi) or copper depletion by overexpression of ATP7 in a cell autonomous manner^13,20,34–37^. We find that inhibition of specific candidate genes enhances ATP7 knockdown phenotypes in epidermal epithelial cells in a sex dependent manner with some genes showing interactions with both sexes while others only interacting in a sex-specific manner. We focused on Lrrk, the fly homologue of Lrrk2 which is the most commonly mutated gene in familial forms of PD and found that it genetically interacts with ATP7 in both males and females in the epidermal epithelium but only in female dopaminergic neurons. These data suggest that disruptions in copper homeostasis may broadly affect genes involved in PD/neurodegeneration and that these interactions may contribute to disease pathology.

## Results

### Coessentiality network analysis of ATP7A interactome

We previously found that loss of ATP7A, a model of increased cellular copper in human cells results in elevated levels of the neurodegeneration associated gene UCHL1 and that UCHL1 inhibition rescues ATP7 phenotypes in human cell culture and flies^38^, suggesting that other neurodegeneration genes in the ATP7A interactome could modulate copper-dependent phenotypes as well. We first tested potential genetic interactions between ATP7A and the interactome PD/neurodegeneration genes by performing a coessentiality network analysis in silico (Figure S1). We assessed genome-scale fitness correlations between ATP7A with ARPC1A, ARPC1B, GIGYF2, HIP1, LRKK2, VPS13C and VPS35 mining the Dependency Map (DepMap) datasets^39,40^. Using the FIREWORKS tool (Fitness Interaction Rank-Extrapolated netWORKs)^41^, we identified that correlated and anticorrelated genes reveal pathways and regulatory mechanisms connecting genes of interest with ATP7A positively (red edges) and negatively (blue edges) interacting with Parkinson’s and neurodegeneration genes. These in silico findings support the model that ATP7A genetic interactions of similar complexity may also occur in mammals as we find in *Drosophila*.

### ATP7 Genetically Interacts with Parkinson’s Disease Genes

We next tested if these predicted interactions from our coessentiality network analysis are biological relevant using our fly ATP7 model (Table 1). In addition, we also tested two other interactome candidates SCYL1 (yata in flies), a neurodegeneration associated protein which had low expression levels, and the PD gene Synj1, which is a component of the endocytic pathway that was identified in the interactome (Table 1)^13,14^. Knockdown of ATP7 in the fly dorsal epidermal epithelium using pnr-GAL4 increases intracellular copper resulting in a decrease in pigmentation in a central stipe along the back of the fly in both females and males due to increased levels of intracellular copper (Figures 1B and 2B) as we and others have previously observed. In addition, these flies also display a widening of the gap between the midline bristles of the thorax (Figures 1B and 2B) as compared to out-crossed control pnr-GAL4 flies (Figures 1A and 2A). We next tested each of the ATP7A interactome genes and find that knockdown of Arpc1 results in disorganized and missing midline bristles in females and males (Figures S2B and S3B). Inhibition of either Gyf or yata results in missing scutellar bristles and some loss of thoracic bristles (Figures S2C, J-L) in females with more mild phenotypes in males (Figures S3C, J-L). Both Synj and Vps35 knockdown also leads to mild loss of scutellar bristles in females and males (Figures S2E, I and S3E, I). We also observe sex differences with inhibition of Hip1 and Vps13 which have no phenotype in females (Figures S2D, G-H) and mild scutellar bristle loss in males (Figures S3D, G-H).

**Table 1.** *Drosophila* Homologues of ATP7A Interactome Genes.

| **Drosophila Candidate Gene** | **Drosophila Gene Abbreviation** | **Human Homolog Gene Name** | **Human Homolog Abbreviation** | **OMIM Number** | **PARK Number** |
| --- | --- | --- | --- | --- | --- |
| Actin-related protein 2/3 complex, subunit 1 | Arpc1 | Actin-Related Protein 2/3 Complex | ARPC1 | 604223 | N/A |
| Gigyf | Gyf | GRB10 interaction GYF protein 2 | GIGYF2 | 607688 | N/A |
| Huntingtin interacting protein 1 | Hip1 | Huntingtin interacting protein 1 | HIP1 | 601767 | PARK11 |
| Leucine-rich repeat kinase | Lrrk | Leucine rich repeat kinase 2 | Lrrk2 | 607060 | PARK8 |
| Synaptojanin | Synj | Synaptojanin 1 | Synj1 | 615530 | PARK20 |
| Vacuolar protein sorting 13 | Vps13 | Vacuolar protein sorting 13C | VPS13C | 616840 | PARK23 |
| Vacuolar protein sorting 35 | Vps35 | VPS35 retromer complex component | VPS35 | 601501 | PARK17 |
| yata | yata | SCYL like pseudokinase 1 | SCYL1 | 616719 | N/A |

**Figure 1.**
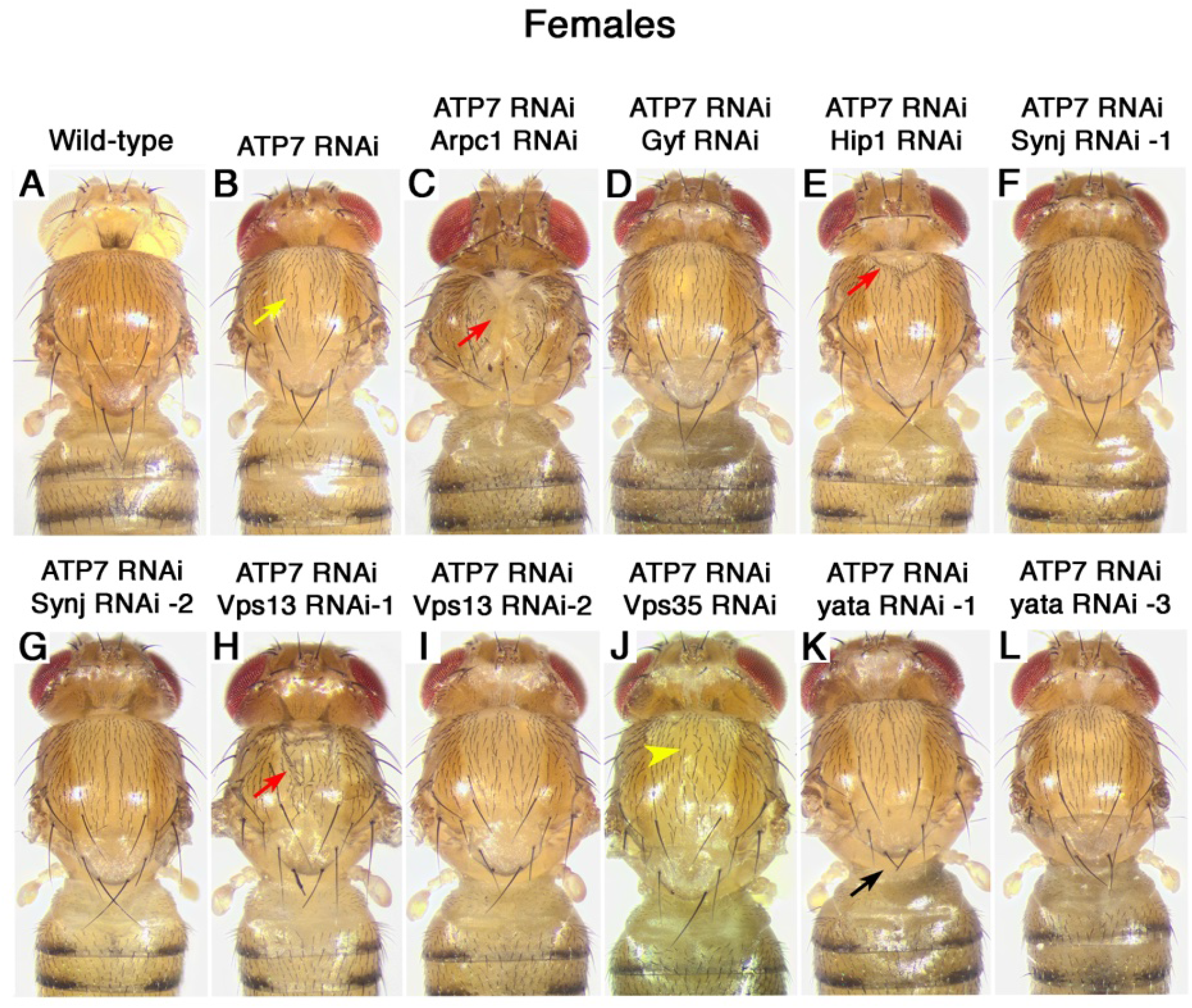
ATP7 Interacts with Parkinson’s Disease/Neurodegeneration Genes in the Epidermal Epithelium in Females. Images of inhibition of ATP7 alone **A)** or in combination with inhibition of different ATP7 interactome homologues using the pnr-GAL4 in females **B-L)**. Yellow arrows indicate widening of the thoracic midline bristles, red arrows indicate areas of thoracic caving in, black arrows show scutellar bristle defects, and yellow arrowhead indicates bristle disorganization.

**Figure 2.**
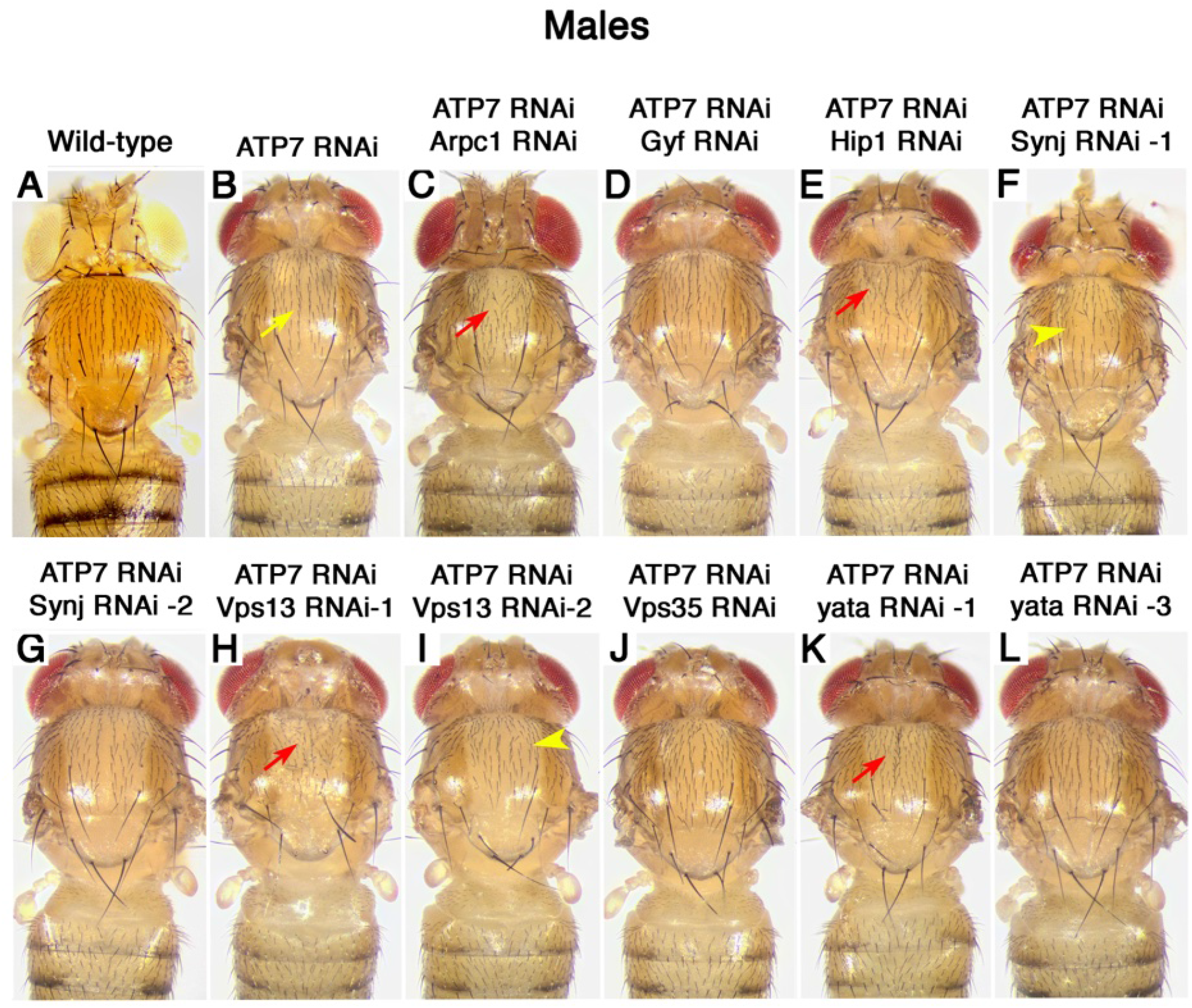
ATP7 Interacts with Parkinson’s Disease/Neurodegeneration Genes in the Epidermal Epithelium in Males. Images of inhibition of ATP7 alone **A)** or in combination with inhibition of different ATP7 interactome homologues using the pnr-GAL4 in males **B-L)**. Yellow arrows indicate areas of thoracic caving in, and red arrows indicate areas of thoracic caving in, black arrows show scutellar bristle defects, and yellow arrowhead indicates missing and disorganized thoracic bristles.

We next tested for genetic interactions between ATP7 down-regulation and inhibition of these interactome genes. We find that in females, inhibition of either Arpc1, Hip1, or Vps13 enhances the ATP7 RNAi phenotype, resulting in a caved in thorax (Figure 1C, E, H, and Table S1). In addition, the ATP7 RNAi phenotype is also enhanced in females with inhibition of yata (line 1) with mild inhibition leading to increased bristle loss (Figure 1K and Table S1) and strong inhibition (line 2) leading to lethality. We also observe an enhancement in disorganization of the midline thoracic bristles when Vps35 was inhibited (Figure 1J and Table S1). However, not all of the interactome genes interacted with ATP7 in females with neither inhibition of Gyf nor Synj affecting the ATP7 down-regulation phenotype (Figure 1D, F-G and Table S1).

As ATP7 is on the X chromosome, males may be more sensitive to genetic perturbations than females, and we find that similar to females Arpc1, Hip1, and Vps13 inhibition enhances the ATP7 thorax caving in phenotype (Figure 2C, E, H and Table S1), though the effect of Arpc1 is not as strong as in females (Figure 2C vs 1C and Table S1). In addition, we find that mild inhibition of yata (line 1) leads to a more severe enhancement of the ATP7 down-regulation phenotype in males as compared to females (Figures 2K vs 1K and Table S1). Though Vps35 inhibition enhanced the ATP7 knockdown phenotype in females, we find no interaction in males (Figure 2J and Table S1). In addition, unlike in females, inhibition of Synj (line 1) also leads to enhancement of the phenotype (Figure 2F and Table S1), whereas similar to females Gyf inhibition does not interact with ATP7 down-regulation in males (Figure 2D and Table S1). These data suggest that normal expression of PD/neurodegeneration genes play a role in protecting cells from increased intracellular copper and that males are more sensitive to these perturbations.

### Inhibition of Lrrk enhances ATP7 down-regulation epidermal epithelium phenotypes

Lrrk2 is the most commonly mutated gene in familial forms of PD^42^ and regulates vesicular trafficking, autophagy, and lysosomal function within cells^17,43–45^. Therefore, we next tested if the Lrrk2 Drosophila orthologue, Lrrk, also plays a role in copper homeostasis using three different RNAi lines (Table S2). We find that in both females and males, knockdown of Lrrk alone does not affect the epidermal epithelium (Figures 3A-D, 4A-D, and Table S1). However, when combined with down-regulation of ATP7, inhibition of Lrrk robustly enhanced the ATP7 RNAi thorax phenotype, yet to a different extent across lines (Figures 3F-H, 4F-H, and Table S1). Knockdown of Lrrk results in a more severe phenotype than in males than in females. For line 1, some males present with necrosis of the scutellum (Figure 4F’) and line 2 displayed a more pronounced caving of the thorax (Figure 4G). In addition, we find that line 3 presented with mild phenotypes in females (Figure 3H and Table S1), whereas males had a more pronounced phenotype with increased thoracic caving (Figure 4H and Table S1) though these phenotypes were milder than line 2 (Figure 4G and Table S1). These data suggest that Lrrk acts to protect epidermal epithelial cells from high levels of intracellular copper and is a more potent interactor in males.

**Figure 3.**
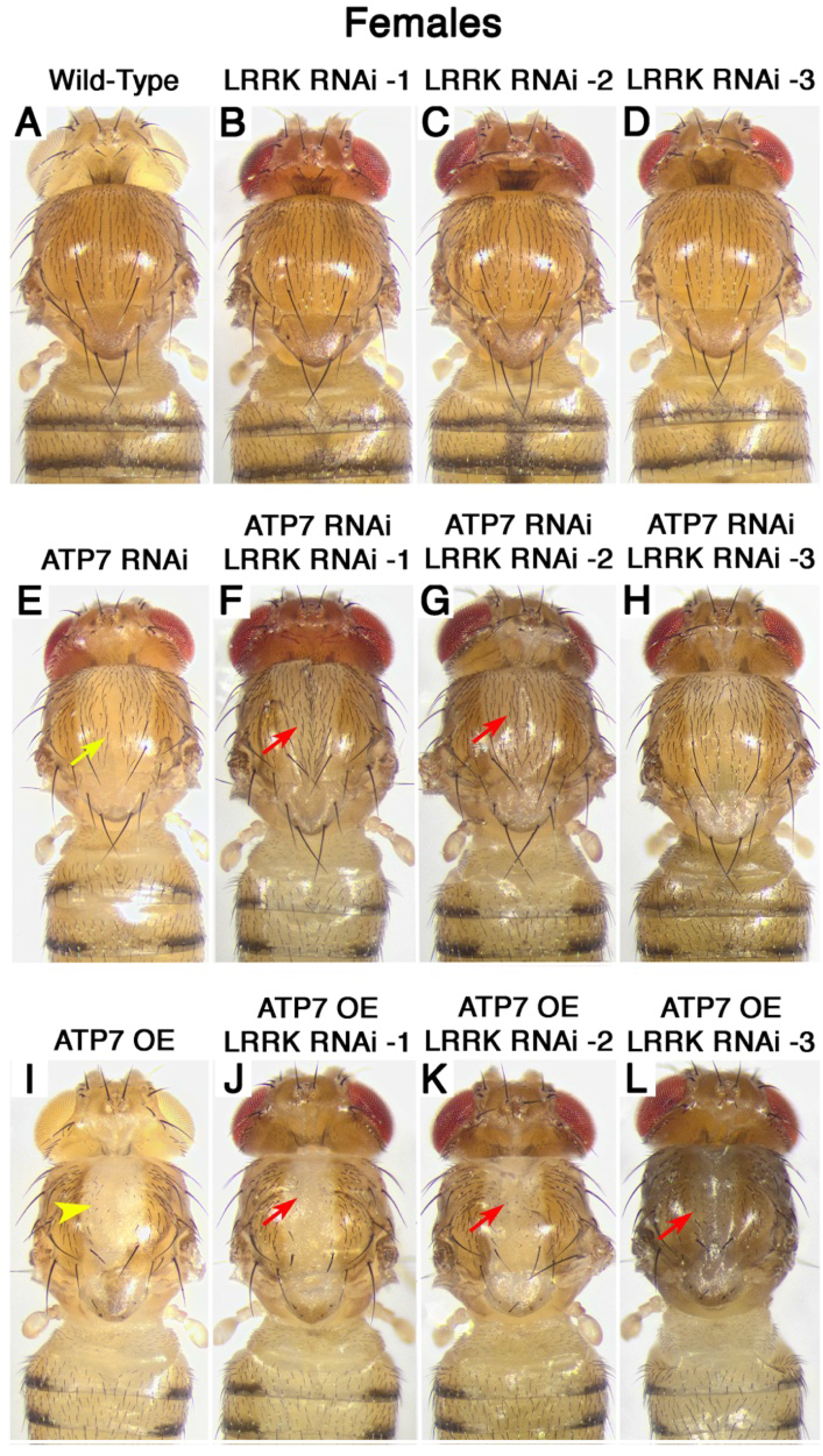
ATP7 Interacts with Lrrk in the Epidermal Epithelium in Females. Images of control **A)** and different Lrrk RNAi lines **B-D)** using the pnr-GAL4 in females. ATP7 down-regulation alone **E)** or in combination with inhibition of Lrrk **F-H)**. ATP7 over-expression **I)** alone or in combination with inhibition of Lrrk **J-L)**. Yellow arrows indicate widening of the bristles along the midline of the thorax. Red arrows indicate thoracic caving in, and yellow arrowheads indicate bristle loss.

**Figure 4.**
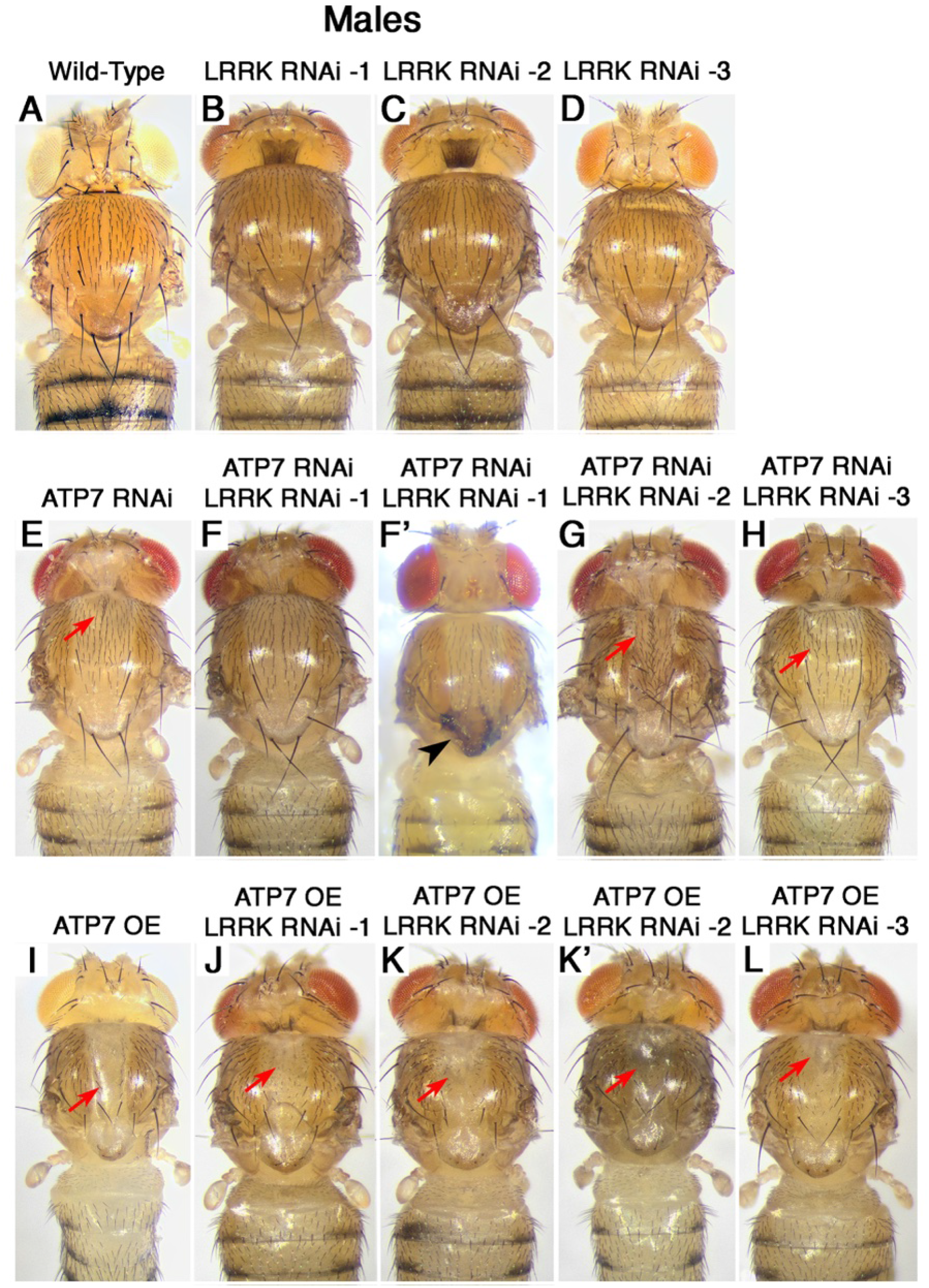
ATP7 Interacts with Lrrk in the Epidermal Epithelium in Males. Images of control **A)** and different Lrrk RNAi lines **B-D)** using the pnr-GAL4 in males. ATP7 down-regulation alone **E)** or in combination with inhibition of Lrrk **F-H)**. ATP7 over-expression **I)** alone or in combination with inhibition of Lrrk **J-L)**. Red arrows indicate thoracic caving in, and black arrowheads indicate necrotic tissue.

### Lrrk and ATP7 down-regulation do not interact to regulate viability

As inhibiting Lrrk leads to a worsening of the ATP7 down-regulation thorax phenotypes, we tested if this alters organismal viability. Though down-regulation of ATP7 does not affect viability in either females or males (Figure S4A, D and Table S3), Lrrk inhibition results in reduced viability (Figure S4B-C, E-F and Table S3). Furthermore, inhibiting Lrrk in the ATP7 down-regulation background reduces viability as compared to ATP7 knockdown alone, though this is not significantly different from Lrrk inhibition alone (Figure S4B-C, E-F and Table S3). These data show that the epithelial damage induced by disruption of the Lrrk and ATP7 interaction spares animal viability.

### Inhibition of Lrrk rescues ATP7 down-regulation dopaminergic neuronal phenotypes in a sex dependent manner

We have previously found that altering copper homeostasis in dopaminergic neurons, which are lost in PD, decreases survival to a copper challenge^13,14^. Therefore, we tested whether Lrrk affects these outcomes. We find that in both females and males, inhibition of either ATP7 or Lrrk alone or in combination does not affect survival on glucose only food (Table S4). Knockdown of ATP7 decreases survival in females when exposed to copper, whereas knockdown of Lrrk did not affect survival as compared to the GAL4 control line (Figure 5A-C, Table S5). However, inhibiting Lrrk in the ATP7 knockdown background, rescued survival (Figure 5A-C, Table S5). In contrast, males died faster when exposed to copper and had decreased survival when either ATP7 or Lrrk is inhibited (Figure 5D-F, Table S5). Knocking down Lrrk in the ATP7 down-regulation background did not significantly alter survival as compared to Lrrk or ATP7 down-regulation (Figure 5D-F, Table S5). These data show that Lrrk contributes to the toxicity caused by increased intracellular copper in females but not males.

**Figure 5.**
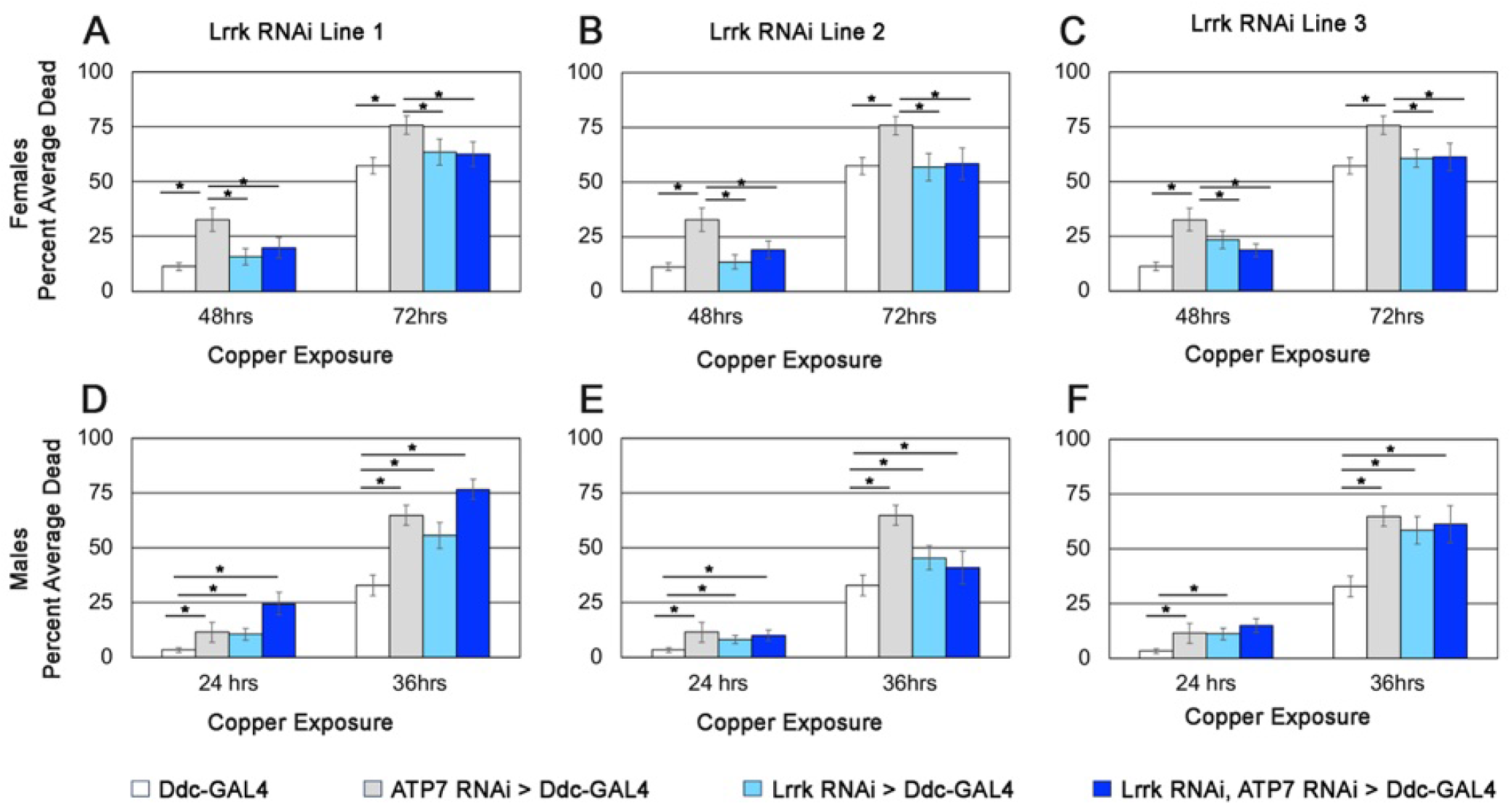
Lrrk Interacts with ATP7 Down-regulation and Copper Exposure in Dopaminergic Neurons. Female **A-C)** and male **D-F)** flies exposed to copper with either inhibition of ATP7 alone or in combination with Lrrk inhibition using the Ddc-GAL4.

### Inhibition of Lrrk enhances ATP7 over-expression epidermal epithelium phenotypes

In humans, both increased intracellular copper and decreased intracellular copper lead to neurological disorders, therefore, we tested if inhibition of Lrrk also interacts with decreased copper levels modelled by over-expression of ATP7. As we and others have previously reported, over-expression of ATP7 in the epidermal epithelium results in decreased pigmentation and bristle loss in both males and females (Figures 3I and 4I and^9,20,34–38^). When we combine inhibition of Lrrk with over-expression of ATP7, we find increased caving in of the thorax (Figures 3J-L, 4J-L, and Table S1) and in some instances a darkening of the thorax (Figure 3L and Figure 4K’) that is indicative of underlying internal necrosis^46,47^. Therefore, inhibition of Lrrk further enhances intracellular copper depletion thorax phenotypes in both females and males. Taken together, our data suggest that Lrrk normally acts to protect against both accumulation (Figures 3E-H, 4E-H, and Table S1) and depletion of intracellular copper (Figures 3I-L, 4I-L, and Table S1).

### Inhibition of Lrrk reduces viability of ATP7 over-expression

We next tested if Lrrk interacts with copper depletion for organismal viability and find that similar to ATP7 down-regulation, over-expression of ATP7 does not affect viability (Figure 6 and Table S6). However, inhibition of Lrrk in the ATP7 over-expression background reduces viability in females and males (Figure 6B-C, E-F and Table S6) but only leads to a more severe reduction as compared to Lrrk inhibition alone in females (Figure 6A-C and Table S6). These data indicate that in females, Lrrk acts to protect epidermal epithelial cells from copper depletion but not accumulation toxicity during development to promote survival to adulthood.

**Figure 6.**
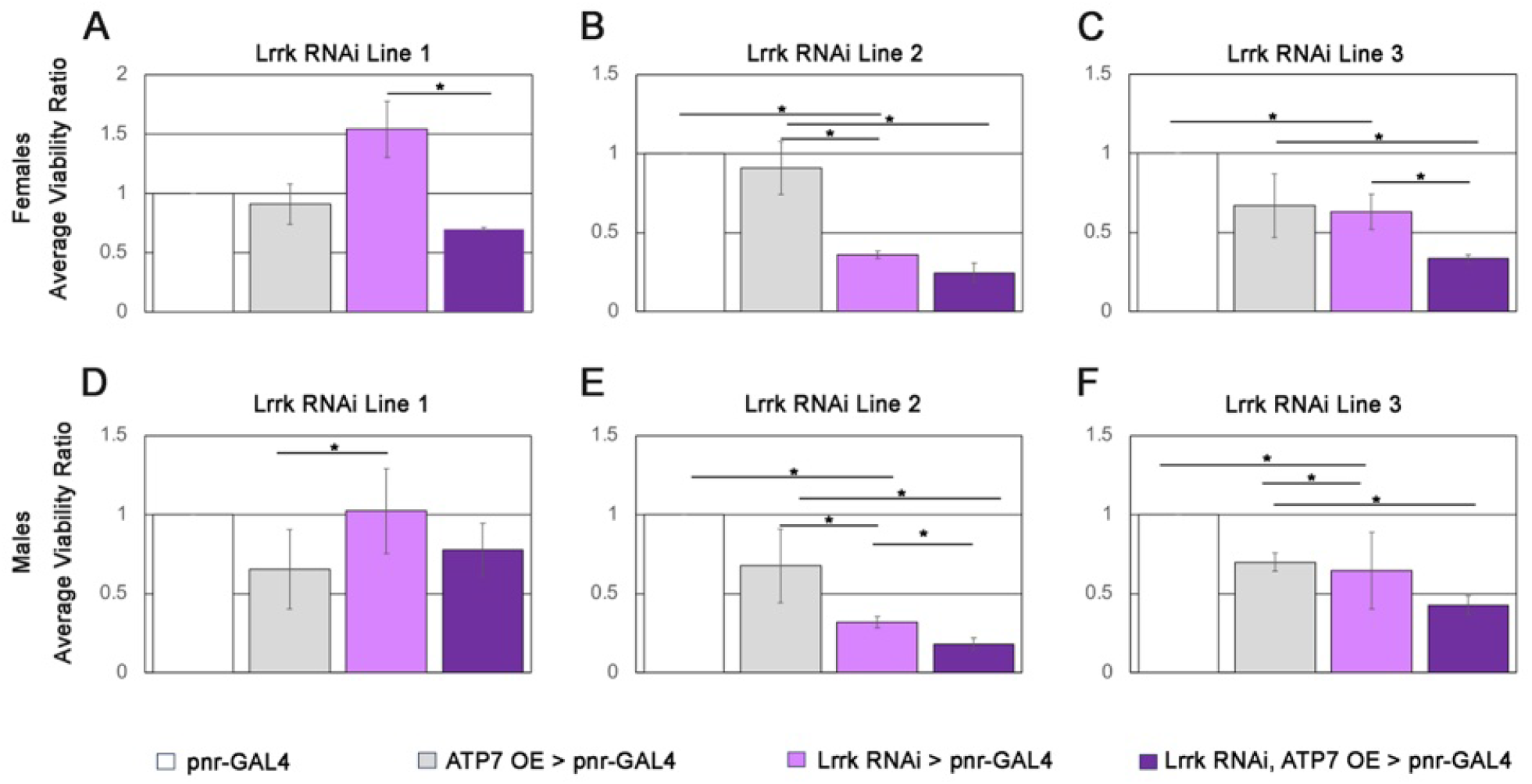
Lrrk Interacts with ATP7 Over-expression and Viability. The normalized ratio of female **A-C)** and male **D-F)** flies that were expected to reach adulthood (viability) with either down-regulation of ATP7 alone or in combination with Lrrk inhibition using the Ddc-GAL4. For Lrrk RNAi line 1 females, statistical results ξ^2^ = 10.93, df = 3, p-value = 0.01211. For Lrrk RNAi line 2 females, statistical results ξ^2^ = 120.44, df = 3, p-value < 2.2e-16. For Lrrk RNAi line 3 females, statistical results ξ^2^ = 41.642, df = 3, p-value = 4.778e-09. For Lrrk RNAi line 1 males, statistical results ξ^2^ = 12.034, df = 3, p-value = 0.007268. For Lrrk RNAi line 2 males, statistical results ξ^2^ = 141.32, df = 3, p-value < 2.2e-16. For Lrrk RNAi line 3 females, statistical results ξ^2^ = 31.117, df = 3, p-value = 8.03e-07.

### Inhibition of Lrrk in dopaminergic neurons reduces survival of ATP7 over-expression in response to environmental copper exposure

As copper depletion in the brain leads to neurological disorders, we next tested if Lrrk interacts with ATP7 over-expression in dopaminergic neurons. In females, over-expression of ATP7 leads to decreased survival, which is not affected by inhibition of Lrrk (Figure 7A-C, Table S7). However, in males ATP7 over-expression had no effect on survival in response to copper exposure though inhibition of Lrrk resulted in decreased survival (Figure 7D-F, Table S7). In addition, Lrrk inhibition in the ATP7 over-expression background also led to decreased survival to levels similar to Lrrk inhibition alone (Figure 7D-F, Table S7), suggesting that inhibition of Lrrk does not sensitize the cell to copper depletion. Therefore, Lrrk does not interact with ATP7 over-expression in dopaminergic neurons in response to environmental copper exposure.

**Figure 7.**
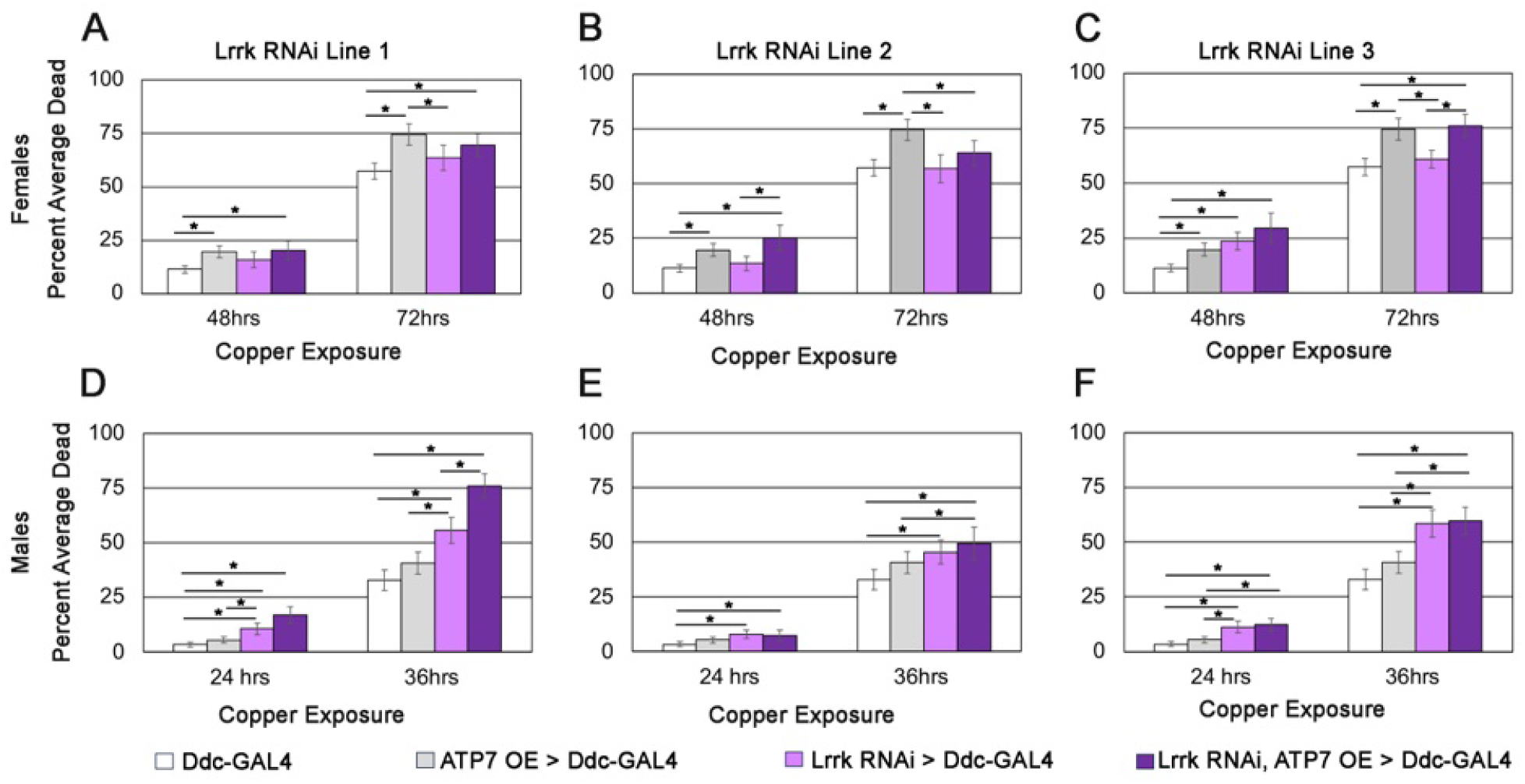
Lrrk Interacts with ATP7 Over-expression and Copper Exposure in Dopaminergic Neurons. Female **A-C)** and male **D-F)** flies exposed to copper with either over-expression of ATP7 alone or in combination with Lrrk inhibition using the Ddc-GAL4.

## Discussion

### Copper homeostasis and PD

Copper dysregulation and PD share many key features including increased oxidative stress, protein aggregation, and mitochondrial damage^21,22^. In addition, PD has been associated with disruptions in copper homeostasis and copper-associated toxicity^21^ with PD associated genes intersecting with signaling pathways that regulate metal homeostasis and counteract oxidative stress^44^. We have previously performed proteomic analyses to identify interactors of the copper transporter ATP7A and identified a number of genes associated with PD and/or neurodegeneration^13,14^. We further confirmed a genetic interaction between ATP7A and one of these candidate genes, UCHL1, which is elevated in ATP7A mutant fibroblasts and inhibition of UCHL1 rescued ATP7A phenotypes^14^ and may be a potential target for the development of new therapeutics. Here, we have further explored how wide-spread this relationship is by testing for genetic interactions between ATP7 and other candidate genes from the ATP7A interactome^13,14^.

We find that a subset (7 out of the 8 genes tested) genetically interact with ATP7 and play a role in protecting epidermal epithelial cells from increased intracellular copper. Five of these genes (Lrrk, Hip1, Synj, Vps13, and Vps35) are associated with PD, while Arpc1 (ARPC1) interacts with known PD genes^48–51^. Two common themes emerge about the functions of those genes that interacted with ATP7: 1) a role in mitochondrial function/health (Vps13, Vps35, and Arpc1)^44,49,50,52–56^ and 2) a role in endocytosis and Golgi trafficking (Vps13, Vps35, Arpc1, yata and Hip1)^4,54,57–60^. In mammalian cell culture, VPS13C plays a role in ER-late endosomes/lysosome communication^57^ and localizes to the outer mitochondrial membrane^53^ where it may also mediate communication between the mitochondria and the ER/endosomal system. Vps13 (VPS13C in mammals) also leads to mitochondrial dysfunction and damage and plays a role in parkin/PINK1 mediated mitophagy^53^. Similarly, a mouse model of VPS35 recapitulated many PD phenotypes including mitochondrial dysfunction and damage^54^ and impaired mitophagy in SH-SY5Y cells^55^. Interestingly both VPS13C and VPS35 play a role in iron homeostasis^61,62^, along with our findings, these data suggest that these genes may play a role in responding to metal dyshomeostasis. Arpc1 regulates actin polymerization and plays a role in a variety of cellular processes including the endosomal trafficking^59^. Upon mitochondrial damage, actin filaments are assembled by Arp2/3 around the damaged mitochondria and inhibition of Arp2/3 impacts mitochondrial fusion^63^ and parkin mediated mitophagy^56^. In addition Arp2/3 is required for the function of the retromer complex, which includes VPS35^51^, suggesting that the nexus between mitochondrial and endosomal-lysosomal processes may be a key target for the copper induced toxicity and PD.

Though HIP1 and yata have no known mitochondrial functions, they also play a role in endocytosis and endosomal trafficking. HIP1 regulates clathrin-mediated endocytosis^60^ and trafficking of oxidative stress responsive cell receptors^64^, and Hip1 loss of function has been implicated in Huntington’s disease, a neurodegenerative disorder in which oxidative stress and mitochondrial dysfunction are prominent pathogenic features^65^, suggesting that HIP1-dependent trafficking may have downstream effects on mitochondrial homeostasis. yata (SCYL1 in mammals) is associated with several neurodegenerative diseases and is an accessory factor for the components of coatomer I (COPI) complex to regulate trafficking between the Golgi and ER^58^. Inhibition of yata in both mice and flies leads to neurodegeneration^66,67^ and interacts with other proteins associated with neurodegenerative diseases such as APPL (Amyloid Precursor Protein-like) in flies^66^ and TDP-43 in mice^68^. We have previously found that ATP7 interacts with the conserved oligomeric Golgi (COG) complex, a Golgi apparatus vesicular tether^20^. This connection between intracellular trafficking and copper homeostasis suggests that when these trafficking genes are disrupted it could exacerbate copper toxicity by impairing copper sequestration, exocytosis, or recycling or metal binding proteins.

### Sex differences in response to copper dysregulation

PD and other neurological disorders can present with skewed sex ratios. For example, men are 1.5 times more likely to develop PD than women^69,70^. In addition, as ATP7A is a sex-linked gene, Menkes disease primarily affects males^71^. Similarly, we find sex-dependent interactions among these genes with Synj interacting with ATP7 in males but not females. We also observed differences in severity of phenotypes across sexes as Arpc1 has milder phenotypes in males than females while yata and Lrrk have more severe phenotypes in males than in females. These data suggest that males and females may have different protective mechanisms against copper dysregulation that intersect with some PD-associated genes but not others.

### Lrrk and ATP7 interactions are context dependent

Lrrk is a kinase that regulates diverse intracellular processes, such as: mitophagy, innate immunity, vesicular transport, and autophagy^17,44,45^, pathways that are frequently disrupted during metal-induced stress, including copper toxicity^20,72–76^.

We find that the interaction between Lrrk and ATP7 are dependent on the context. For example, we find that Lrrk protects against both increased and depleted levels of intracellular copper in epidermal epithelial cells to maintain thoracic structure in both sexes. However, in response to increased copper, males are more severely affected when Lrrk is inhibited. Furthermore, only in females and only during intracellular copper depletion in the epidermal epithelium does Lrrk act to promote organismal viability. These interactions are also cell type specific as we find that inhibition of Lrrk in females, but not males, improved survival in response to increased levels of intracellular copper (ATP7 down-regulation with copper exposure) but had no effect on survival in response to depletion of intracellular copper (ATP7 over-expression with copper exposure). These data suggest that Lrrk contributes to intracellular copper toxicity, and since it plays a role in stress response pathways^17^, Lrrk2 may help buffer cells against copper-induced toxicity. Additionally, as the pathogenic G2019S mutation in Lrrk is a gain of function mutation^77,78^, disruptions in copper homeostasis could lead to an earlier age of onset or more severe symptoms/faster disease progression.

Through this study, we have confirmed that the ATP7A interactome identified from human cell culture and patient samples^13,38^, represent biologically significant interactions that contribute to how cells respond to disruptions in cellular homeostasis. In addition, we also find that there is an intimate and common connection between copper homeostasis and genetic forms of PD, which may also provide new insights into how environmental exposure to copper and other heavy metals contribute to sporadic forms of PD.

## Materials and Methods

### Coessentiality Analysis

Coessentiality Analysis was performed with the FIREWORKS engine^41^ using ATP7A with ARPC1A, ARPC1B, GIGYF2, HIP1, LRKK2, VPS13C and VPS35 genes as entries in the Pan-Cancer dataset as Context. The engine was run with the top 5 primary nodes and the top five secondary nodes positively and negatively correlated.

### Drosophila Strains and Husbandry

The following strains were obtained from the Bloomington *Drosophila* Stock Center, Bloomington, IL: w1118 (#5905), Ddc-Gal4 (#7009), pnr-Gal4 (#3039), UAS-Lrrk-RNAi-1 (#32457), UAS-Lrrk-RNAi-2 (#39019), and UAS-Lrrk-RNAi-3 (#35249). The UAS-ATP7-RNAi (#108159) stock was obtained from the Vienna *Drosophila* Resource Center. The UAS-ATP7-wt (overexpression) was a gift from Richard Burke, Monash University, Australia. See Table S2 for a full list of genotypes. All stocks were reared on standard Molasses Food (Genesee Scientific) at 25°C in 12hr:12hr light:dark cycle.

### Imaging

ATP7A interactome candidate genes were knocked down in epidermal epithelium cells in either an ATP7 loss of function or overexpression background using the pnr-Gal4 driver. 1-week-old flies were imaged using a LEICA S6D stereomicroscope with a Leica EC4 Digital Camera. Z-stacks were then aligned and max projections generated using Zerene Stacker.

### Viability Assay

Three biological replicates of 8-10 virgin females were crossed with 4-5 males for each genotype. The flies were transferred into new vials every three days resulting in three technical replicates for each cross. Once the offspring began eclosing, they were tracked for nine days, accounting for the number of males and females for each expected genotype were recorded. All crosses were reared on standard Molasses Food (Genesee Scientific) at 25°C in 12hr:12hr light:dark cycle.

### Copper Toxicity Assay

Copper feedings were performed on 1-week-old virgin females and males reared separately. ATP7 was either knockdown or overexpressed alone or in combination with Lrrk knockdown in the dopaminergic neurons using the Ddc-Gal4 driver. Flies were starved for 3 hours and transferred to empty vials with Whatman glass microfiber filter discs saturated in either 1 mM CuSO4 + 5% glucose solution or 5% glucose solution. Survival was assayed for every 12-24 hours for five days. Experiments were performed with a minimum of 10 replicates per genotype per sex with ∼10 flies per replicate at 25°C in 12hr:12hr light:dark cycle.

### Statistical Analysis

The results of the copper toxicity assay were tested for statistical significance testing survival and proportion data. Survival data was tested using Kaplan-Meier survival analysis, log-rank tests, and pairwise comparisons to assess the significance between genotypes and survival time. Proportion data for copper survival and viability were tested by creating a generalized linear model and an analysis of deviance table was used to fit variance using chi-squared tests for significance, per-time contrasts were conducted with post emmeans testing with tukey adjustment. All statistical analyses were performed in R.

## Acknowledgements

We would also like to acknowledge the Bloomington Drosophila Stock Center (NIH P40OD018537) and the Vienna Drosophila Resource Center for providing fly stocks used in this study. Funding for this project included start-up funds from Illinois State University and Oregon State University, and R15AR070505 to AVM, and 1RF1AG060285 to VF. EW is supported by NIH grant ES034796 and NG is supported by 1T32GM160704-01. The content is solely the responsibility of the authors and does not necessarily represent the official views of the National Institutes of Health. The funders had no role in study design, data collection and analysis, decision to publish, or preparation of the manuscript.

## Author Contributions

BA and NG performed experiments and data analysis. EW, VF, and AVM made intellectual contributions to experimental design, data analysis and interpretation, and provided funding. All authors were involved in reading and editing the manuscript.

## Data Availability Statement

The data that support the findings of this study are available from the corresponding author upon reasonable request.

