## Supplemental data for "Context-dependent ATP7 Interactions with Parkinson’s Disease-associated Genes Modulate Copper Homeostasis Phenotypes"

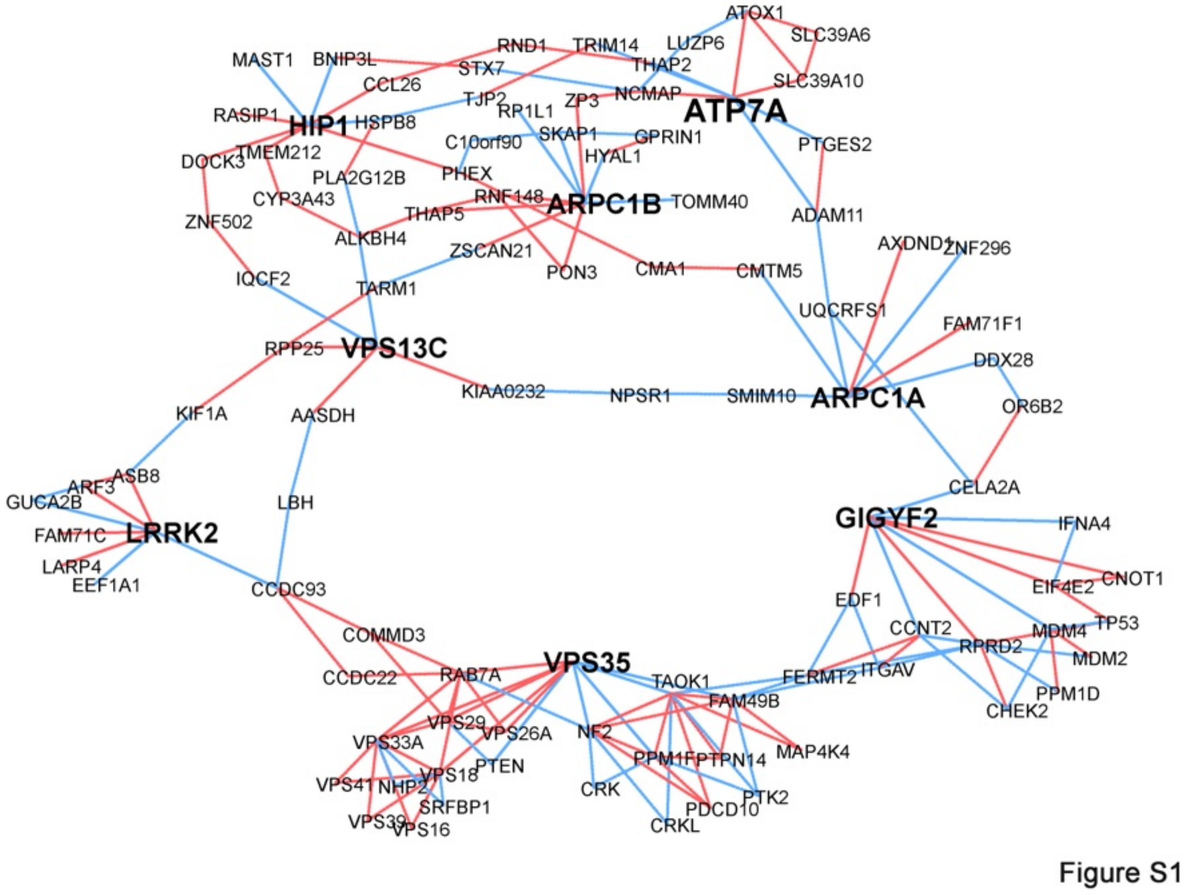


**Figure S1. Coessentiality Network Analysis of ATP7A.**

Coessentiality network analysis showing genome-scale fitness correlations of ATP7A and Parkinson’s and neurodegeneration genes as inputs. Nodes correspond to first and second level interaction nodes. Network of correlated (red edges) and anticorrelated genes (blue edges) was built with the FIREWORKS tool.

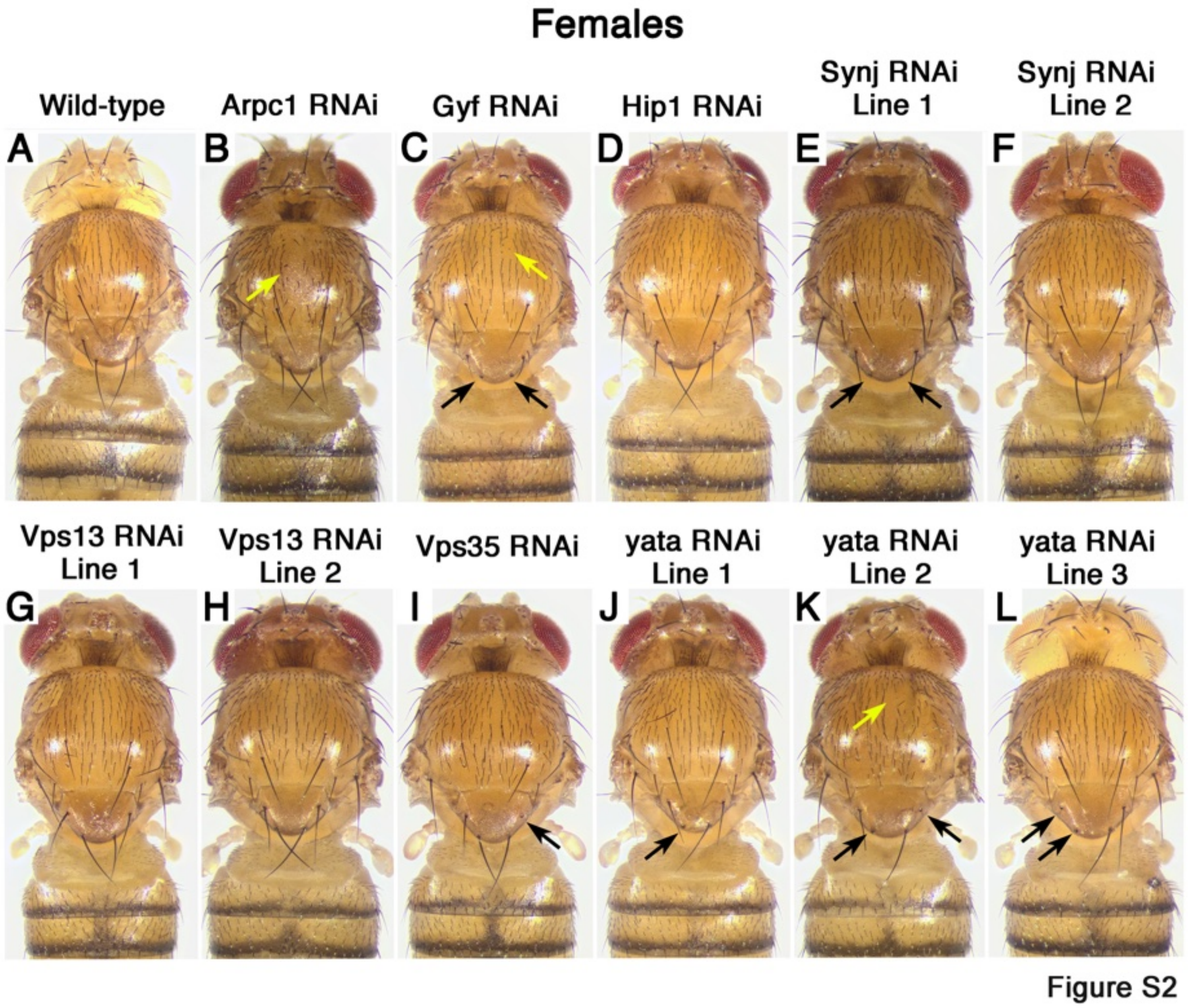


**Figure S2. Inhibition of Lrrk in the Epidermal Epithelium in Females.** Images of inhibition of ATP7 interactome genes using the pnr-GAL4 in females. Yellow arrows indicate missing and disorganization thoracic midline bristles. Black arrows show loss of scutellar bristles.

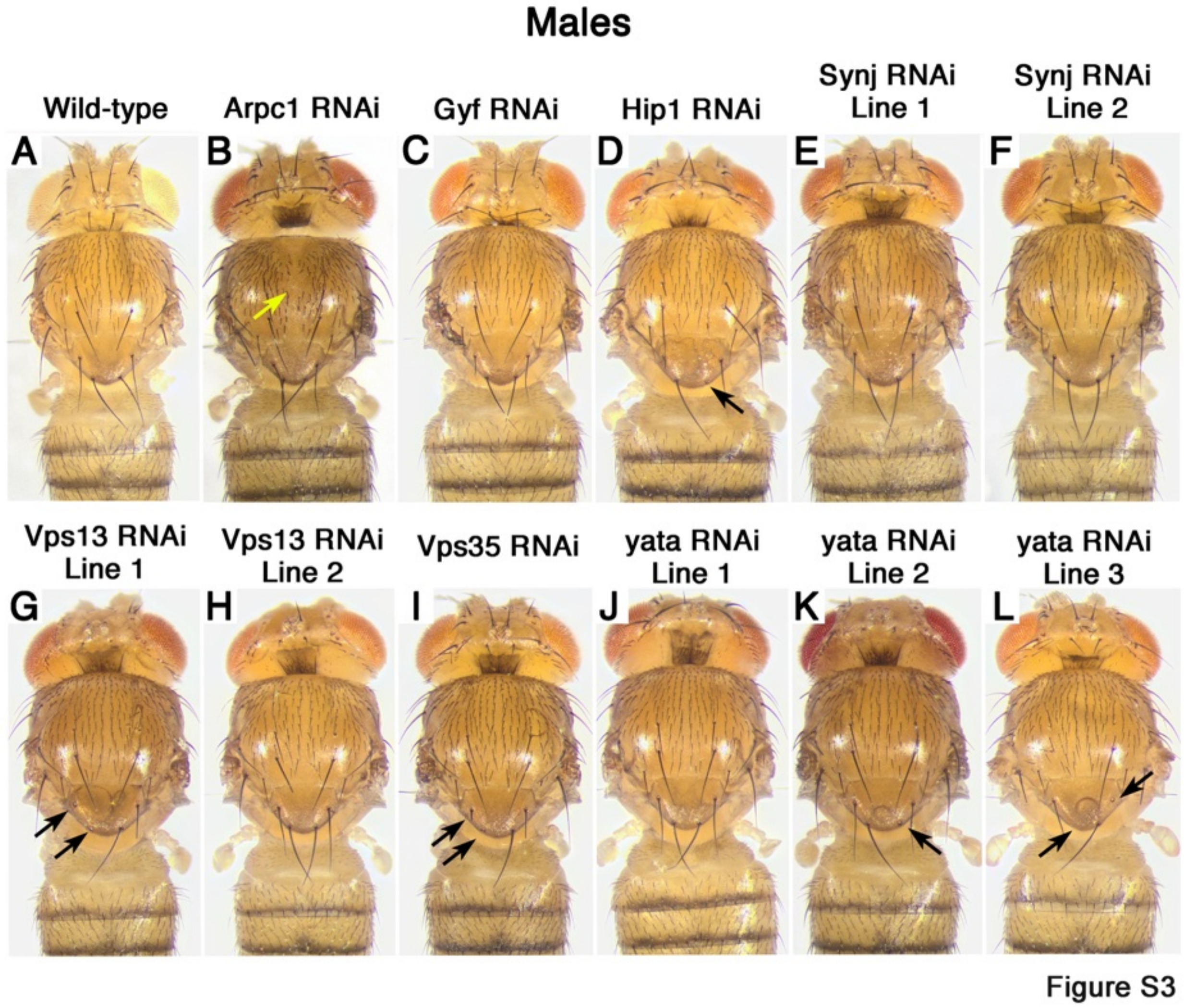


**Figure S3. Inhibition of Lrrk in the Epidermal Epithelium in Males.** Images of inhibition of ATP7 interactome genes using the pnr-GAL4 in females. Yellow arrows indicate missing and disorganization thoracic midline bristles. Black arrows show loss of scutellar bristles.

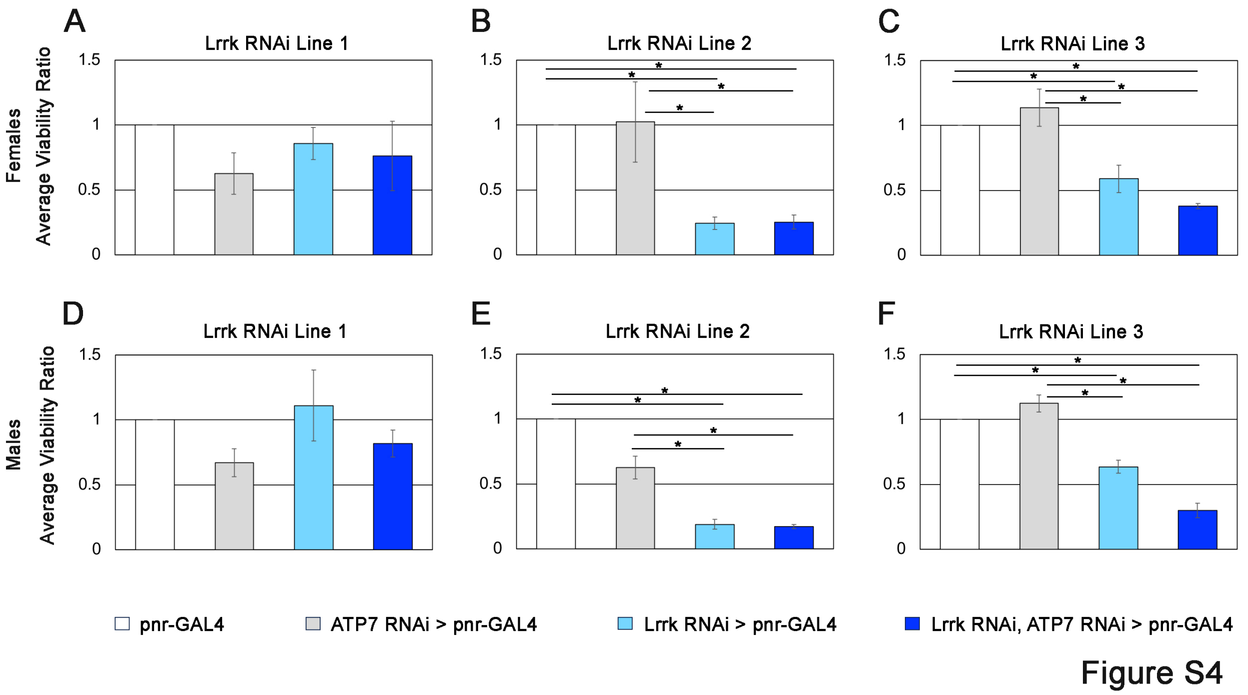


**Figure S4. Lrrk Interacts with ATP7 Down-regulation and Viability.** The normalized ratio of female **A-C) and** male **D-F)** flies that were expected to reach adulthood (viability) with either down-regulation of ATP7 alone or in combination with Lrrk inhibition using the Ddc-GAL4. For Lrrk RNAi line 1 females, statistical results $\chi^2 = 7.1566$, df = 3, p-value = 0.06707. For Lrrk RNAi line 2 females, statistical results $\chi^2 = 112.05$, df = 3, p-value < 2.2e-16. For Lrrk RNAi line 3 females, statistical results $\chi^2 = 47.872$, df = 3, p-value = 2.267e-10. For Lrrk RNAi line 1 males, statistical results $\chi^2 = 6.052$, df = 3, p-value = 0.1091. For Lrrk RNAi line 2 males, statistical results $\chi^2 = 97.556$, df = 3, p-value < 2.2e-16. For Lrrk RNAi line 3 females, statistical results $\chi^2 = 64.697$, df = 3, p-value = 5.823e-14.

| Gene | ATP7 knockdown Interaction in Females | ATP7 knockdown Interaction in Males |
|---|---|---|
| Arpc1 | enhancement | milder enhancement than females |
| Gyf | no interaction | no interaction |
| Hip1 | enhancement | enhancement |
| Synj line 1 | no interaction | enhancement |
| Synj line 2 | no interaction | no interaction |
| Vps13 line 1 | enhancement | enhancement |
| Vps13 line 2 | mild enhancement | mild enhancement |
| Vps35 | enhancement | no interaction |
| yata line 1 | mild enhancement | stronger enahncement than females |
| yata line 2 | severe enhancement (lethality) | severe enhancement (lethality) |
| yata line 3 | no interaction | no interaction |
| Lrrk Line 1 | enhancement | stronger enahncement than females |
| Lrrk Line 2 | enhancement | stronger enahncement than females |
| Lrrk Line 3 | mild enhancement | stronger enahncement than females |
| | **ATP7 OE Interaction in Females** | **ATP7 OE Interaction in Males** |
| Lrrk Line 1 | enhancement | enhancement |
| Lrrk Line 2 | enhancement | enhancement |
| Lrrk Line 3 | enhancement | enhancement |

**Table S1. Summary of Interactions Between ATP7 Down-regulation and Interactome Genes in the Epidermal Epithelium**

| Stock Number | Genotype | Short Name |
|---|---|---|
| BDSC 5905 | w[1118] | $w^{1118}$ |
| BDSC 7009 | w[1118]; P{w[+mC]=Ddc-GAL4.L}Lmpt[4.36] | Ddc-Gal4 |
| BDSC 3039 | y[1] w[1118]; P{w[+mW.hs]=GawB}pnr[MD237]/TM3, P{w[+mC]=UAS- y.C}MC2, Ser[1] | pnr-Gal4 |
| VDRC 108159 | P{KK100396}VIE-260B | UAS-ATP7 RNAi |
| Gift of R. Burke | UAS-ATP7-wt | UAS-ATP7-wt |
| BDSC 31246 | y[1] v[1]; P{y[+t7.7] v[+t1.8]=TRiP.JF01763}attP2 | UAS-Arpc1 RNAi |
| BDSC 28896 | y[1] v[1]; P{y[+t7.7] v[+t1.8]=TRiP.HM05106}attP2 | UAS-Gyf RNAi |
| BDSC 32504 | y[1] sc[*] v[1] sev[21]; P{y[+t7.7] v[+t1.8]=TRiP.HMS00508}attP2 | UAS-Hip1 RNAi Line 1 |
| BDSC 27489 | y[1] v[1]; P{y[+t7.7] v[+t1.8]=TRiP.JF02639}attP2 | UAS-Synj RNAi Line 1 |
| BDSC 34378 | y[1] sc[*] v[1] sev[21]; P{y[+t7.7] v[+t1.8]=TRiP.HMS01368}attP2 | UAS-Synj RNAi Line 2 |
| BDSC 42625 | y[1] sc[*] v[1] sev[21]; P{y[+t7.7] v[+t1.8]=TRiP.HMS02460}attP40 | UAS-Vps13 RNAi Line 1 |
| BDSC 38270 | y[1] sc[*] v[1] sev[21]; P{y[+t7.7] v[+t1.8]=TRiP.HMS01715}attP40 | UAS-Vps13 RNAi Line 2 |
| BDSC 38944 | y[1] sc[*] v[1] sev[21]; P{y[+t7.7] v[+t1.8]=TRiP.HMS01858}attP40 | UAS-Vps35 RNAi |
| BDSC 32986 | y[1] sc[*] v[1] sev[21]; P{y[+t7.7] v[+t1.8]=TRiP.HMS00786}attP2 | UAS-yata RNAi Lline 1 |
| VDRC 110214 | P{KK102668}VIE-260B | UAS-yata RNAi Line 2 |
| VDRC 19275 | w1118; P{GD8890}v19275 | UAS-yata RNAi Line 3 |
| BDSC 32457 | y[1] sc[*] v[1] sev[21]; P{y[+t7.7] v[+t1.8]=TRiP.HMS00456}attP2 | UAS-Lrrk RNAi Line 1 |
| BDSC 39019 | y[1] v[1]; P{y[+t7.7] v[+t1.8]=TRiP.HMS01937}attP40 | UAS-Lrrk RNAi Line 2 |
| BDSC 35249 | y[1] sc[*] v[1] sev[21]; P{y[+t7.7] v[+t1.8]=TRiP.GL00136}attP2/TM3, Sb[1] | UAS-Lrrk-RNAi Line 3 |

**Table S2.** ***Drosophila*** **Genotypes**

| | Females | Males |
|---|---|---|
| Lrrk RNAi Line 1 | viability | viability |
| pnr-GAL4 vs Lrrk RNAi | 0.6411 | 0.4637 |
| pnr-GAL4 vs ATP7 RNAi | 0.0654 | 0.7236 |
| pnr-GAL4 vs Lrrk RNAi and ATP7 RNAi | 0.1915 | 0.9989 |
| Lrrk RNAi vs ATP7 RNAi | 0.4799 | 0.0693 |
| Lrrk RNAi vs Lrrk RNAi and ATP7 RNAi | 0.7966 | 0.6288 |
| ATP7 RNAi vs Lrrk RNAi and ATP7 RNAi | 0.9575 | 0.6874 |
| Lrrk RNAi Line 2 | | |
| pnr-GAL4 vs Lrrk RNAi | <.0001 | <.0001 |
| pnr-GAL4 vs ATP7 RNAi | 0.6794 | 0.4638 |
| pnr-GAL4 vs Lrrk RNAi and ATP7 RNAi | <.0001 | <.0001 |
| Lrrk RNAi vs ATP7 RNAi | <.0001 | <.0001 |
| Lrrk RNAi vs Lrrk RNAi and ATP7 RNAi | 0.9991 | 0.7364 |
| ATP7 RNAi vs Lrrk RNAi and ATP7 RNAi | <.0001 | <.0001 |
| Lrrk RNAi Line 3 | | |
| pnr-GAL4 vs Lrrk RNAi | 0.0025 | <.0001 |
| pnr-GAL4 vs ATP7 RNAi | 0.8549 | 0.9995 |
| pnr-GAL4 vs Lrrk RNAi and ATP7 RNAi | <.0001 | <.0001 |
| Lrrk RNAi vs ATP7 RNAi | 0.0002 | <.0001 |
| Lrrk RNAi vs Lrrk RNAi and ATP7 RNAi | 0.2072 | 0.4632 |
| ATP7 RNAi vs Lrrk RNAi and ATP7 RNAi | <.0001 | <.0001 |

**Table S3. Statistical Comparisons of Lrrk Inhibition and ATP7 Down-regulation Viability**

| **ATP7 Down-regulation** | **Glucose Treatment** | | | | **ATP7 Over-expression** | **Glucose Treatment** | | |
|---|---|---|---|---|---|---|---|---|
| | Precent Average Flies Dead | | | | | Precent Average Flies Dead | | |
| Females | 24hrs | 48hrs | 72hrs | | Females | 24hrs | 48hrs | 72hrs |
| w1118 Ddc-GAL4 | 0 | 0 | 0 | | w1118 Ddc-GAL4 | 0 | 0 | 0 |
| ATP7 RNAi Ddc-GAL4 | 0 | 0 | 0 | | ATP7 wt Ddc-GAL4 | 0 | 0 | 0 |
| Lrrk 1 Ddc-GAL4 | 0 | 0 | 0 | | Lrrk 1 Ddc-GAL4 | 0 | 0 | 0 |
| Lrrk 1 ATP7 RNAi Ddc-GAL4 | 0 | 0 | 0.595 | | Lrrk 1 ATP7 wt Ddc-GAL4 | 0 | 0.556 | 0.556 |
| Lrrk 2 Ddc-GAL4 | 0 | 0 | 0 | | Lrrk 2 Ddc-GAL4 | 0 | 0 | 0 |
| Lrrk 2 ATP7 RNAi Ddc-GAL4 | 0 | 0 | 0.955 | | Lrrk 2 ATP7 wt Ddc-GAL4 | 0 | 0.455 | 0.909 |
| Lrrk 3 Ddc-GAL4 | 0 | 0 | 0.357 | | Lrrk 3 Ddc-GAL4 | 0 | 0 | 0.357 |
| Lrrk 3 ATP7 RNAi Ddc-GAL4 | 0 | 0 | 0 | | Lrrk 3 ATP7 wt Ddc-GAL4 | 0 | 0 | 0 |
| Males | 24hrs | 36hrs | 48hrs | | Males | 24hrs | 36hrs | 48hrs |
| w1118 Ddc-GAL4 | 0 | 0 | 0 | | w1118 Ddc-GAL4 | 0 | 0 | 0 |
| ATP7 RNAi Ddc-GAL4 | 0 | 0 | 0 | | ATP7 wt Ddc-GAL4 | 0 | 0 | 0 |
| Lrrk 1 Ddc-GAL4 | 0.385 | 0.385 | 0.385 | | Lrrk 1 Ddc-GAL4 | 0.385 | 0.385 | 0.385 |
| Lrrk 1 ATP7 RNAi Ddc-GAL4 | 0.455 | 0.455 | 0.455 | | Lrrk 1 ATP7 wt Ddc-GAL4 | 0 | 0 | 0 |
| Lrrk 2 Ddc-GAL4 | 0 | 0 | 0 | | Lrrk 2 Ddc-GAL4 | 0 | 0 | 0 |
| Lrrk 2 ATP7 RNAi Ddc-GAL4 | 0.417 | 1.250 | 1.250 | | Lrrk 2 ATP7 wt Ddc-GAL4 | 0 | 0 | 0 |
| Lrrk 3 Ddc-GAL4 | 0 | 0 | 0 | | Lrrk 3 Ddc-GAL4 | 0 | 0 | 0 |
| Lrrk 3 ATP7 RNAi Ddc-GAL4 | 0 | 1.455 | 1.455 | | Lrrk 3 ATP7 wt Ddc-GAL4 | 1.270 | 1.270 | 1.270 |

**Table S4. Statistical Comparisons of Glucose Treated Lrrk and ATP7 Down-regulation or Over-expression Survival**

| | Females | | Males | |
|---|---|---|---|---|
| Lrrk RNAi Line 1 | 48hrs | 72hrs | 24hrs | 36hrs |
| Ddc-GAL4 vs Lrrk RNAi | 0.3009 | 0.1089 | 0.0002 | <.0001 |
| Ddc-GAL4 vs ATP7 RNAi | <.0001 | p <.0001 | 0.0004 | <.0001 |
| Ddc-GAL4 vs Lrrk RNAi and ATP7 RNAi | 0.056 | 0.5238 | <.0001 | <.0001 |
| Lrrk RNAi vs ATP7 RNAi | <.0001 | 0.0056 | 1.0000 | 0.4022 |
| Lrrk RNAi vs Lrrk RNAi and ATP7 RNAi | 0.7503 | 0.9554 | 0.0001 | 0.0005 |
| ATP7 RNAi vs Lrrk RNAi and ATP7 RNAi | 0.0068 | 0.0041 | 0.0005 | 0.0848 |
| Lrrk RNAi Line 2 | | | | |
| Ddc-GAL4 vs Lrrk RNAi | 0.7917 | 0.9948 | 0.013 | 0.0037 |
| Ddc-GAL4 vs ATP7 RNAi | <.0001 | <.0001 | 0.0004 | <.0001 |
| Ddc-GAL4 vs Lrrk RNAi and ATP7 RNAi | 0.0419 | 0.9072 | 0.0014 | 0.1613 |
| Lrrk RNAi vs ATP7 RNAi | <.0001 | <.0001 | 0.625 | <.0001 |
| Lrrk RNAi vs Lrrk RNAi and ATP7 RNAi | 0.3029 | 0.972 | 0.8299 | 0.7809 |
| ATP7 RNAi vs Lrrk RNAi and ATP7 RNAi | 0.0014 | 0.0001 | 0.9893 | <.0001 |
| Lrrk RNAi Line 3 | | | | |
| Ddc-GAL4 vs Lrrk RNAi | 0.0001 | 0.961 | 0.0001 | <.0001 |
| Ddc-GAL4 vs ATP7 RNAi | <.0001 | <.0001 | 0.0004 | <.0001 |
| Ddc-GAL4 vs Lrrk RNAi and ATP7 RNAi | 0.1344 | 0.6298 | <.0001 | <.0001 |
| Lrrk RNAi vs ATP7 RNAi | 0.0302 | <.0001 | 0.9976 | 0.1569 |
| Lrrk RNAi vs Lrrk RNAi and ATP7 RNAi | 0.784 | 0.8303 | 0.5103 | 0.9352 |
| ATP7 RNAi vs Lrrk RNAi and ATP7 RNAi | 0.0269 | 0.0162 | 0.4584 | 0.6078 |

**Table S5. Statistical Comparisons of Copper Treated Lrrk and ATP7 Down-regulation Survival**

| | Females | Males |
|---|---|---|
| Lrrk RNAi Line 1 | viability | viability |
| pnr-GAL4 vs Lrrk RNAi | 0.6934 | 0.6924 |
| pnr-GAL4 vs ATP7 OE | 0.9921 | 0.0634 |
| pnr-GAL4 vs Lrrk RNAi and ATP7 OE | 0.1148 | 0.9616 |
| Lrrk RNAi vs ATP7 OE | 0.5481 | 0.0057 |
| Lrrk RNAi vs Lrrk RNAi and ATP7 OE | 0.0075 | 0.445 |
| ATP7 OE vs Lrrk RNAi and ATP7 OE | 0.2436 | 0.2395 |
| Lrrk RNAi Line 2 | | |
| pnr-GAL4 vs Lrrk RNAi | <.0001 | <.0001 |
| pnr-GAL4 vs ATP7 OE | 0.9921 | 0.1037 |
| pnr-GAL4 vs Lrrk RNAi and ATP7 OE | <.0001 | <.0001 |
| Lrrk RNAi vs ATP7 OE | <.0001 | <.0001 |
| Lrrk RNAi vs Lrrk RNAi and ATP7 OE | 0.0514 | 0.012 |
| ATP7 OE vs Lrrk RNAi and ATP7 OE | <.0001 | <.0001 |
| Lrrk RNAi Line 3 | | |
| pnr-GAL4 vs Lrrk RNAi | 0.009 | <.0001 |
| pnr-GAL4 vs ATP7 OE | 0.0523 | 0.1208 |
| pnr-GAL4 vs Lrrk RNAi and ATP7 OE | <.0001 | 0.0002 |
| Lrrk RNAi vs ATP7 OE | 0.8967 | 0.0377 |
| Lrrk RNAi vs Lrrk RNAi and ATP7 OE | 0.003 | 0.9634 |
| ATP7 OE vs Lrrk RNAi and ATP7 OE | 0.0002 | 0.042 |

**Table S6. Statistical Comparisons of Lrrk Inhibition and ATP7 Over-expression Viability**

| | Females | | Males | |
|---|---|---|---|---|
| Lrrk RNAi Line 1 | 48hrs | 72hrs | 24hrs | 36hrs |
| Ddc-GAL4 vs Lrrk RNAi | 0.3009 | 0.1089 | 0.0002 | <.0001 |
| Ddc-GAL4 vs ATP7 OE | 0.0088 | <.0001 | 0.3108 | 0.2611 |
| Ddc-GAL4 vs Lrrk RNAi and ATP7 OE | 0.0187 | 0.0034 | <.0001 | <.0001 |
| Lrrk RNAi vs ATP7 OE | 0.4831 | 0.0274 | 0.0357 | <.0001 |
| Lrrk RNAi vs Lrrk RNAi and ATP7 OE | 0.5551 | 0.4744 | 0.384 | 0.0019 |
| ATP7 OE vs Lrrk RNAi and ATP7 OE | 1.0000 | 0.693 | 0.0002 | <.0001 |
| Lrrk RNAi Line 2 | | | | |
| Ddc-GAL4 vs Lrrk RNAi | 0.7917 | 0.9948 | 0.013 | 0.0037 |
| Ddc-GAL4 vs ATP7 OE | 0.0088 | <.0001 | 0.3108 | 0.2611 |
| Ddc-GAL4 vs Lrrk RNAi and ATP7 OE | 0.0001 | 0.3282 | 0.05 | <.0001 |
| Lrrk RNAi vs ATP7 OE | 0.1192 | <.0001 | 0.4784 | 0.3608 |
| Lrrk RNAi vs Lrrk RNAi and ATP7 OE | 0.0031 | 0.4792 | 0.9877 | 0.4629 |
| ATP7 OE vs Lrrk RNAi and ATP7 OE | 0.5691 | 0.0109 | 0.7451 | 0.0135 |
| Lrrk RNAi Line 3 | | | | |
| Ddc-GAL4 vs Lrrk RNAi | 0.0001 | 0.961 | 0.0001 | <.0001 |
| Ddc-GAL4 vs ATP7 OE | 0.0088 | <.0001 | 0.3108 | 0.2611 |
| Ddc-GAL4 vs Lrrk RNAi and ATP7 OE | <.0001 | <.0001 | 0.0001 | <.0001 |
| Lrrk RNAi vs ATP7 OE | 0.7487 | <.0001 | 0.0182 | <.0001 |
| Lrrk RNAi vs Lrrk RNAi and ATP7 OE | 0.4599 | 0.0003 | 0.9986 | 0.9988 |
| ATP7 OE vs Lrrk RNAi and ATP7 OE | 0.0978 | 0.9999 | 0.0184 | <.0001 |

**Table S7. Statistical Comparisons of Copper Treated Lrrk and ATP7 Over-expression Survival**
